# Assaying *Chlamydia pneumoniae* persistence in monocyte-derived macrophages identifies schisandrin lignans as phenotypic switchers

**DOI:** 10.1101/708487

**Authors:** Eveliina Taavitsainen, Maarit Kortesoja, Leena Hanski

## Abstract

Antibiotic-tolerant persister bacteria involve frequent treatment failures, relapsing infections and the need for extended antibiotic treatment. Taking persisters into account in susceptibility assays is thus an essential success factor in antibacterial drug discovery. The virulence of the obligate intracellular bacterium *Chlamydia pneumoniae* is tightly linked to its propensity for persistence, but current susceptibility screening on this gram-negative respiratory pathogen relies on permissive epithelial cells. To establish an improved antichlamydial susceptibility assay allowing the analysis of both actively growing and persister bacteria, we studied *C. pneumoniae* clinical isolate CV-6 infection kinetics in THP-1 macrophages by qPCR and quantitative culture. Indicated by the steady increase of chlamydial genome copy numbers and infectious progeny as well as the failure of azithromycin to eradicate the intracellular forms of the bacterium, the macrophages were found to harbor a subpopulation of persister *C. pneumoniae* cells. The potential of the assay for the discovery of anti-persister molecules against intracellular bacteria was demonstrated by the identification of the differential effects of two dibenzocyclooctadiene lignans on *C. pneumoniae* infection. While schisandrin reverted *C. pneumoniae* persistence and promoted productive infection, schisandrin C was superior to azithromycin in eradicating the *C. pneumoniae* infection. The phenotypic switch was associated with the suppression of cellular glutathione pools, implying that targeting glutathione homeostasis may provide a novel means for intracellular bacteria resuscitation. In conclusion, these data highlight the value of macrophages over permissive cell lines in anti-persister agent discovery on intracellular bacteria and targeting host cell redox status to fight persistent infections.

## Background

In the course of evolution, bacteria have developed various means for protecting themselves from unfavorable conditions. Described as a reversible dormant phenotype, persistence has been acknowledged as one major survival strategy of bacteria^1^. Bacterial persistence is considered major cause of antibiotic treatment failures and relapsing infections and it also contributes to the rise of antibiotic resistance ^2^. Owing to redundant mechanisms, these phenotypical variants are able to survive under antibiotic pressure and revert back to metabolically more active phenotype when stressful conditions are cleared off.

In clinical settings, bacterial dormancy is associated with hard-to-treat infections via two mechanistically overlapping phenomena, persistent infections evading host immune responses and antibiotic persistence defined based on the presence of drug-tolerant subpopulations of bacteria. Both of these features are typical to infections caused by *Chlamydia pneumoniae*, a gram-negative obligate intracellular human pathogen that causes respiratory infections from dry cough to pneumonia. While a majority of *C. pneumoniae* infections are subclinical, nearly everyone getting infected during their lifetime, the bacterium is also responsible for 5-10% of community-acquired pneumonia cases worldwide ^3, 4^. *C. pneumoniae* has a unique biphasic development cycle, where the bacteria switch between an infectious form elementary body (EB) and a non-infectious metabolically active reticulate body (RB) ^5^. The acute phase may also be followed by a persistent infection ^6^, occurring spontaneously in monocytes and macrophages ^4, 7^. A morphological hallmark of the persistent phenotype is the emergence of abnormal reticulate bodies with low metabolic activity and replication ^6, 8^. However, cellular and molecular mechanisms driving chlamydial persistent infections are considered heterogeneous, redundant and not yet fully understood ^9, 10^.

Besides the acute respiratory illnesses, *C. pneumoniae* has been related to many chronic inflammatory diseases, such as atherosclerosis and asthma exacerbation ^11, 12^. The ability of *C. pneumoniae* to persist in infected cell populations forms the basis for the hypotheses on these disease connections, and monocytes and macrophages have a main role in the initiation of the chlamydial persistence ^4, 6^.

Both *in vitro* and *in vivo*, the outcome of *C. pneumoniae* infection in monocytes and macrophages depends on the bacterial strain and host cell origin ^4, 13, 14^. *C. pneumoniae* is able to infect and survive inside alveolar and PBMC-derived macrophages ^13, 15, 16^. In these cells, *C. pneumoniae* has been described to form small and non-mature or persistent-like inclusions and produce not at all or significantly lower yields of infectious progenies than in permissive epithelial cells. ^7^. However, the previous studies reporting limited replication of *C. pneumoniae* in macrophage models, detection of replication has been made solely by quantitative culture ^17, 18^. These studies may reliably evaluate the presence of a productive infection but limitation in detection method leaves intracellular replication and persistent forms of the bacterium beyond notice.

To date, a variety of anti-persister molecules have been described against both gram-positive and gram-negative human pathogens, major strategies involving direct eradication of the metabolically quiescent cells by eg. Membrane-active compounds, bacterial resuscitation by boosting energy metabolism and application of combination therapies ^2^. According to current consensus, taking persister bacteria into account is a critical success factor in antibacterial drug discovery and this paradigm shift has brought stationary phase cultures, bacterial biofilms and other persister-enriched culture systems as essential tools in this respect. Despite such progress, susceptibility testing and antibacterial discovery platforms for intracellular bacteria still mostly rely on conventional methodology and models on actively replicating bacteria. Regarding *C. pneumoniae*, only a few membrane-active agents capable of affecting EB infectivity independent of metabolic activity have been described ^19, 20^ and to date, no agents capable of affecting the persistent intracellular forms of the bacterium have been described. Furthermore, current standard methods for antichlamydial susceptibility testing are based solely on permissive epithelial cells and involve the suppression of host cell responses by cycloheximide treatment ^21, 22^.

To establish an antichlamydial susceptibility assay allowing the simultaneous analysis of actively growing and persistent bacteria, we studied *C. pneumoniae* infection in THP-1 macrophages. The model was found to harbor a mixed infection, involving simultaneously present populations of replicating and persister bacteria and its potential in the discovery of antichlamydial agents targeting also persistent bacteria was demonstrated by the identification of schisandrin and schisandrin C, two dibenzocyclooctadiene lignans with antichlamydial activities that are qualitatively distinct from each other and the reference antibiotic azithromycin.

## Materials and Methods

### Compounds

Schisandrin was obtained from Sigma-Aldrich, St. Louis, MO, USA and schisandrin B and schisandrin C were purchased from Fine Tech Industries, London, UK. The lignans, as well as azithromycin (BioWhittaker, Lonza, Basel, Switzerland) used as a reference antibiotic were dissolved in dimethyl sulfoxide (DMSO) and diluted in cell culture media at indicated concentrations.

### Cell culture

All cell cultures were maintained at 37 °C, 5 % CO_2_ and 95 % air humidity.

THP-1 cells (ATCC TIB-202) were maintained in RPMI 1640 Dutch edition medium (Gibco, Invitrogen, Thermo Fisher Scientific, Massachusetts, USA) supplemented with 10 % FBS (BioWhittaker, Lonza, Basel, Switzerland), 2 mM L-glutamine (BioWhittaker, Lonza, Basel, Switzerland), 0.05 mM merkaptoethanol (Gibco, Invitrogen, Thermo Fisher) and 20 μg/ml gentamicin (Sigma-Aldrich, St. Louis, MO, USA). For differentiation into macrophage-like cells, THP-1 cells were incubated for 48 – 72 h with 0.16 μg/ml `phorbol-12-myristate-13-acetate` (PMA, Sigma-Aldrich, St. Louis, MO, USA). Human HL cells ^23^ were maintained in RPMI 1640 (BioWhittaker, Lonza, Basel, Switzerland) supplemented with 7.5 % FBS, 2 mM L-glutamine and 20 μg/ml gentamicin. When seeding HL cells into well plates, an overnight incubation was applied prior to the experiment.

### Infections

For qPCR and infectious progeny experiments, The cells were seeded into 24-well plates (THP-1 monocytes and HL cells at a density of 4 × 10^5^ cell per well, THP-1 macrophages 3.5 × 10^5^ cell per well) and infected with *C. pneumoniae* (strain CV-6, obtained from professor Matthias Maass, Paracelsus Medical University, Salzburg, Austria, propagated as previously described ^24^). Cell monolayers were centrifuged at 550g for 1 h and incubated 1 h in 37 °C. Then, fresh medium or medium with compounds was added and the cultures were incubated from 24 to 144 h. To determine effect of GSH on C. *pneumoniae* infection, 2 mM GSH ethyl ester was added to the infected cultures at 2, 24 or 48 h post infection. For HL cell infections, cell culture medium was supplemented with 1 μg/ml of cycloheximide (CHX, Sigma-Aldrich, St. Louis, MO, USA).

### Quantitative PCR

DNA from cell cultures was extracted with a GeneJet Genomic DNA purification kit (Thermo Fisher Scientific, Massachusetts, USA) according to the manufacturer’s instructions for mammalian cells. The DNA concentration in samples was measured with Multiskan Sky Microplate spectrophotometer by μDrop Plate and the DNA was stored at −20 °C until use. An established qPCR method on *C. pneumoniae* ompA gene ^25^ was applied to quantify *C. pneumoniae* genome copy numbers. Using Step One plus Real-Time PCR system (Thermo Fisher Scientific, Massachusetts,USA). The primers were selected for qPCR run as follows: forward primer, VD_4_F (5’-TCC GCA as TTG CTC AGC C-3’) and reverse primer, VD_4_R (5’-AAA CAA TTT GCA TGA AGT CTG AGA A-3’). The reactions were prepared in 96-well MicroAmp optical plate by adding 20 ng of extracted DNA to 10 μl of master mix to 20 μl qPRC reaction. Detection was performed with Step One plus Real-Time PCR system by using the manufacturer’s standard protocol. Conditions in thermal cycle were 95 °C for 20 s and 40 cycles of 95 °C for 3 s and 60 °C for 30 s.

### Infectious progeny assay

The EBs were harvested from infected THP-1 macrophages at various time points (72 – 144 h) post infection by collecting the culture supernatants, centrifuging them at 21 000 rpm for 1 h in 4 °C and resuspending the pellets with 0.2 ml of fresh cold media. Monolayer cells from the same samples were scraped to 0.2 ml of fresh cold media. All samples were stored in −80 °C.

The bacterial progeny in the samples was quantified by inoculating them on HL monolayers with CHX. After infection, fresh medium with CHX was added and cells were incubated for 70 h. Then, cells were fixed and stained with a genus-specific anti-LPS antibody (Pathfinder, Bio-rad, California, USA). Chlamydial titers were determined based on the inclusion counts observed with fluorescence microscope.

### Intracellular ROS detection assay

THP-1 macrophages in 96-well plate (6 × 10^4^ cells/well) were subjected to an infection by *C. pneumoniae*, treatment with the schisandrin lignans or a combination of these two, using a culture medium without merkaptoethanol. In experiments with 1 – 4 h exposure, the cells were preloaded with 20 μM DCFH-DA (Sigma-Aldrich, St. Louis, MO, USA) for 30 min and washed with PBS prior to lignan administration. After 24 – 72 h exposures, cells were washed once with PBS, loaded with DCFH-DA for 30 min, washed with PBS and incubated for further 3 hours. After that, fluorescence was recorded at 503/523 nm with Varioskan Lux plate reader.

### Glutathione quantification assay

The intracellular GSH levels of THP-1 macrophages after *C. pneumoniae* infection, lignan treatment or combination of these two were determined using enzymatic recycling method described previously by Rahman et al ^26^.

For data normalization, total protein concentration determination of the cell lysates was performed with acetone precipitation. 100 μl of cell lysate sample was heated 5 min at 95 °C and 400 μl of cold (−20 °C) acetone was added. Sample was mixed and incubated 1 h at −20 °C, centrifuged at 15000g and supernatant was discarded. Pellet was resuspended to 100 mM Tris-buffer (pH; 7.5) and protein concentration was detected with Multiskan sky, μDrop plate. Sample purity was evaluated with 260/280 ratio values.

### Data analysis

Statistical tests were performed using SPSS Statistics 24 software. Differences between means were calculated with Student’s t-test with Bonferroni correction. P values < 0.05 were considered statistically significant. Outliers were defined from data by Grupps test, in significance level 0.05.

## Results and discussion

### THP-1 macrophages as a model for C. pneumoniae persistence

Despite the common use of THP-1 cells as infection hosts in studies on *C. pneumoniae* biology, no comprehensive data on *C. pneumoniae* replication or infection kinetics in these cells have been available. To evaluate the usefulness of this cell line for establishing a model for chlamydial persistence, the growth kinetics of *C. pneumoniae* in both monocytic and PMA-differentiated, macrophage-like, THP-1 cells was followed with qPCR. For comparison, epithelial HL cells hosting an active *C. pneumoniae* infection were included in the study.

Consistent with earlier studies reporting active and efficient replication of *C. pneumoniae* in HL cells ^27^, our data with HL cells demonstrate a continuous increase in the number of chlamydial genome equivalents (GE) detectable since 32 h post infection (Fig. 1A). As expected, treatment with 20 nM azithromycin resulted in 99 % reduction of bacterial genome copy number in these cells.

**Figure 1.**
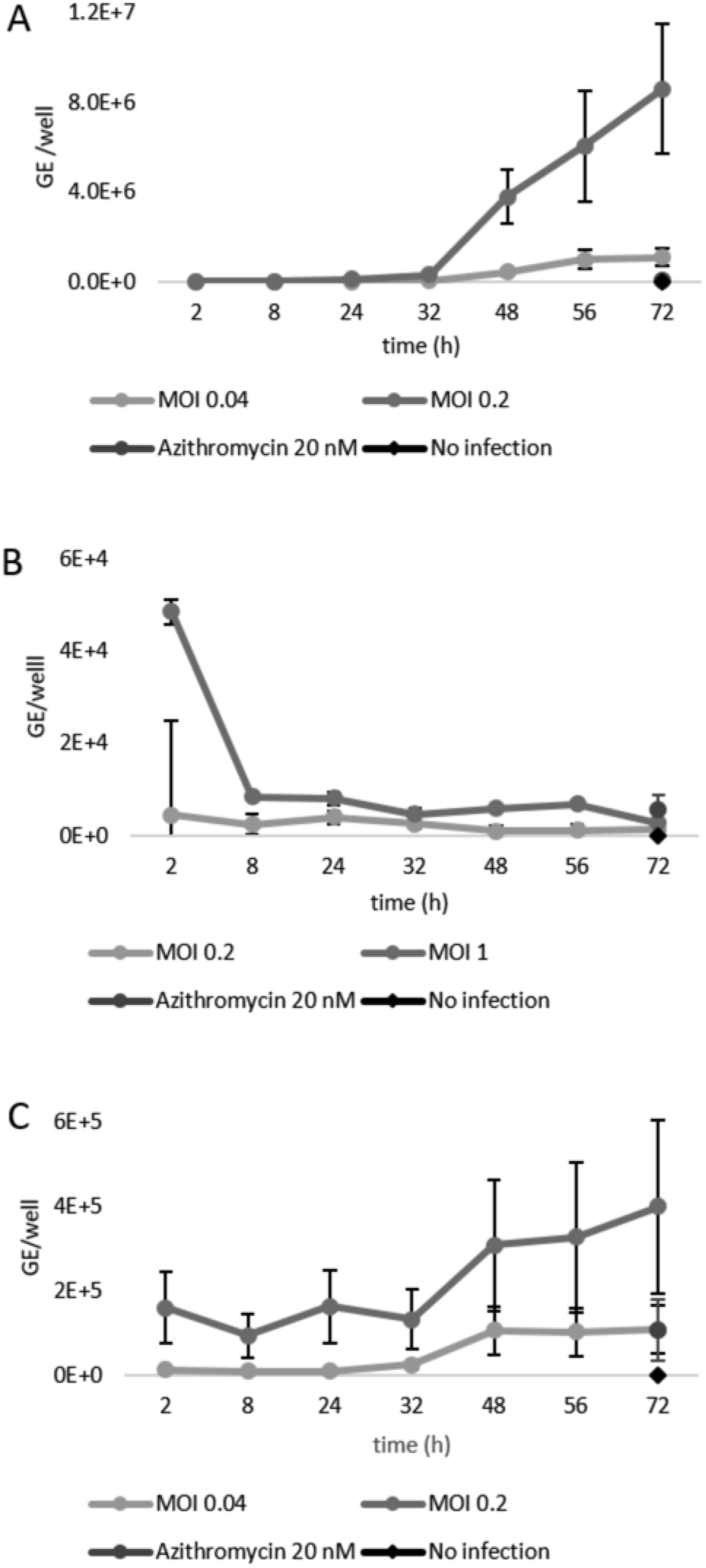
Growth kinetics of *C. pneumoniae* in A) HL epithelial cell B) THP-1 monocytes and C) THP-1 macrophages. Cells were infected at MOI 0.04, 0.2 or 1 IFU/cell and *C. pneumoniae* genome copy numbers were determined with qPCR on ompA gene in seven time points. Data are presented as number of genome equivalents per well and shown as mean ± SEM, n=4. Abbreviations: Genome equivalents, GE, multiplicity of infection, MOI

In monocytic THP-1 cells, the *C. pneumoniae* GE numbers decreased drastically shortly after infection and after 8 h, stayed in a low yet detectable level throughout the observation period (Fig. 1B). Azithromycin (20 nM) had no detectable effect on the chlamydial genome copy numbers.

In contrast to monocytic cells, the GE numbers in infected THP-1 macrophages stayed in constant level until 32 h post infection and increased thereafter throughout the observation time (Fig. 1C). However, *C. pneumoniae* replication in THP-1 macrophages was less efficient than in HL cells since the genome copy numbers in the macrophages were approximately 20-fold lower than in HL cells at 72 h post infection. Treatment with 20 nM azithromycin resulted in only 73 % reduction of GE numbers in THP-1 macrophages. As the qPCR data indicated the presence of actively dividing population of bacterial cells in THP-1 macrophages, we next evaluated whether *C. pneumoniae* is able to establish a productive infection yielding new infectious progeny in these cells.

Within its productive life cycle, *C. pneumoniae* infectious progeny production occurs typically 48 – 72 h post infection, involving differentiation of the newly formed bacterial cells into EBs that leave the host cell to infect neighboring cells. For detecting the production of infectious EB progenies in THP-1 macrophages, subculturing of infected THP-1 cell lysates and culture medium supernatants in HL cells was used.

In these experiments, production of *C. pneumoniae* EBs capable of infecting HL cells was detected in all THP-1 cell lysates and supernatants collected from 72 to 144 h post infection (Table 1), indicating a continuous, asynchronous production of infectious progeny. In general, significantly higher quantities of IFU were observed in cell lysates than in supernatant samples. A duplication in IFU amounts in supernatants and triplication in cell lysates was observed from 72 to 144 h post infection.

**Table 1.** Quantification of infectious *C. pneumoniae* progeny EBs following an infection of THP-1 macrophages.

|  | cell lysate | supernatant |
| --- | --- | --- |
|  | (IFU/ml) | (IFU/ml) |
| 72 h | 44 700 ± 0 | 11 200 ± 1 100 |
| 96 h | 57 700 ± 700 | 11 200 ± 1 100 |
| 120 h | 201 200 ± 20 900 | 18 400 ± 1 800 |
| 144 h | 330 000 ± 27 300 | 21 300 ± 4 000 |

Note. Data are presented as representative examples of IFU/ml values, shown as mean±SEM, n=4. Abbreviations: infectious units per ml, IFU/ml

Based on these data, it was evident that THP-1 macrophages can harbor a productive C. *pneumoniae* infection. However, the limited effectiveness of azithromycin in inhibiting *C. pneumoniae* growth indicates that the infection in this cell population is refractory to this macrolide antibiotic commonly used as a golden standard for treating chlamydial infections and highlights the importance of characterizing antichlamydial compounds also in macrophage-like cells.

### Schisandrin lignans as modulators of C. pneumoniae infection

We have recently identified the antichlamydial activity of dibenzocyclooctadiene lignans isolated from a medicinal plant *Schisandra chinensis* against the actively replicating bacteria in respiratory epithelial cells ^28, 29^. Within the validation process on the current assay, three of these lignans, schisandrin, schisandrin B and schisandrin C were evaluated for their efficacy against *C. pneumoniae* in the THP-1 macrophage model at concentrations determined based on cell viability assays (Supplementary Tables 1 and 2).

Impact of the schisandrin lignans on *C. pneumoniae* growth kinetics was determined in THP-1 macrophages by qPCR in various time points from 2 to 144 h post infection. Of note, azithromycin at its typically used concentration 20 nM did not have any effect against the MOI1 infection (data not shown). At 100 nM azithromycin reduced the *C. pneumoniae* genome numbers by 68 % of infection 72 h post infection and by 93 % at 144 h post infection (Fig. 2), thus showing extended time kill characteristics typical to persistent infections.

**Figure 2.**
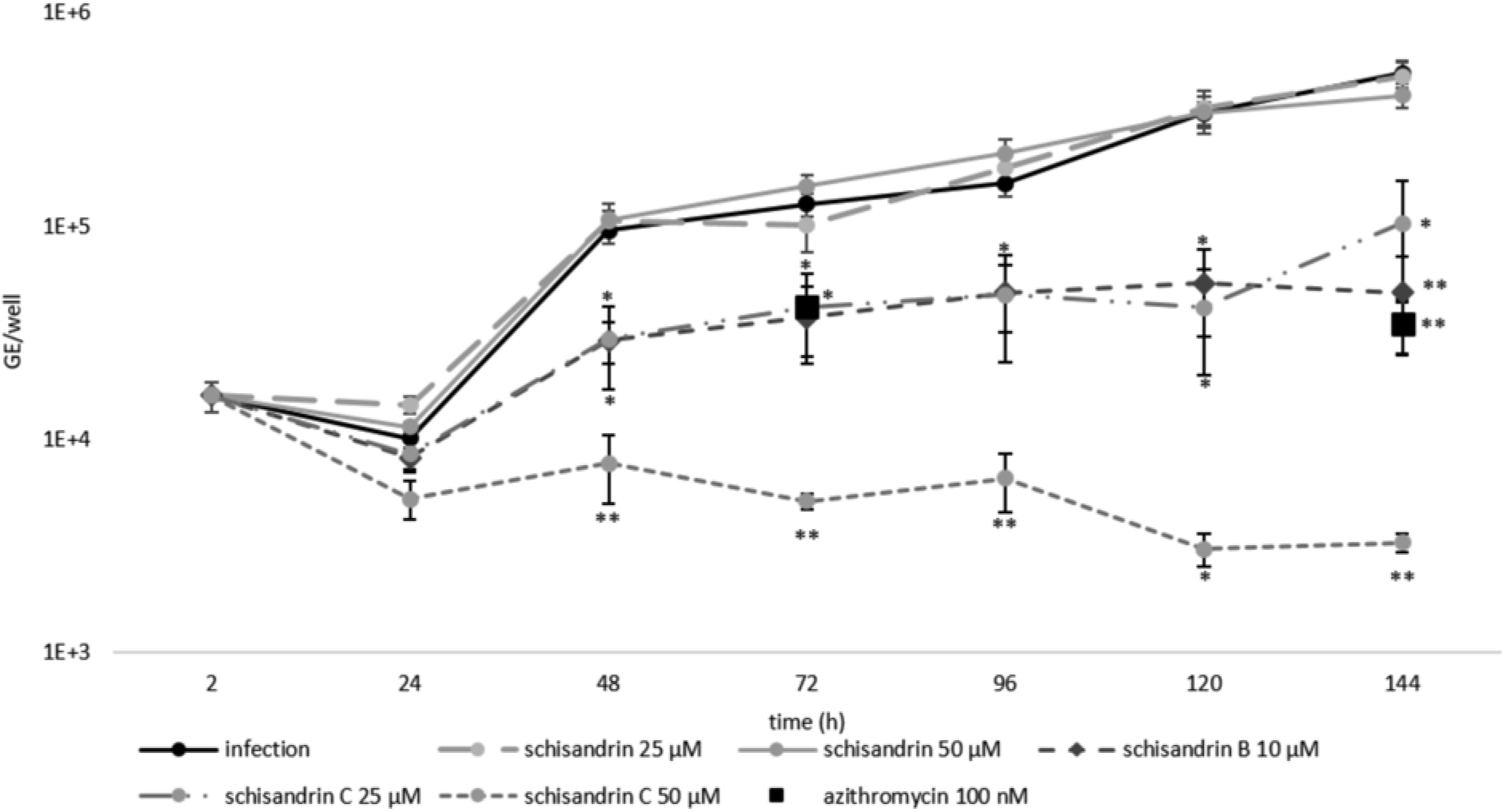
Impact of schisandrin lignans on *C. pneumoniae* growth kinetics in THP-1 macrophages. Cells were infected at MOI 1 IFU/cell and *C. pneumoniae* genome copy numbers were determined with qPCR on ompA gene. Fresh medium was added at 72 h. Data are shown as total genome numbers of *C. pneumoniae* per well ± SEM. Statistical significance is presented as marks of P values: < 0.05: *; < 0.01: **; < 0.001: ***, n=4. Abbreviation: Genome equivalents, GE

As shown in Fig. 2, 10 μM schisandrin B and 25 μM schisandrin C were as effective as 100 nM azithromycin in reducing *C. pneumoniae* genome numbers at 72 h and 144 h after infection. 50 μM schisandrin C was superior to all other samples assayed for *C. pneumoniae* genome number reduction and showed a statistically significant difference to azithromycin in this respect, with 95% reduction in bacterial genome numbers at 72 h and 99 % reduction at 144 h.

The effect of the schisandrin lignans on *C. pneumoniae* infectious progeny production in THP-1 macrophages was also evaluated. As shown in Table 2, schisandrin B and schisandrin C decreased infectious progeny production in a statistically significant manner at 72 and 144 h. In schisandrin C treated samples (25 μM or 50 μM), not any characteristical inclusions were detected, only some small irregular inclusion-like structures were observed in HL monolayers inoculated with the cell lysates and none in those inoculated with supernatant samples. At 50 μM, schisandrin did not have effect on infectious EB production, neither in cell lysates nor in supernatant samples. In contrast, at 25 μM schisandrin treatment resulted in a statistically significant increase in infectious EB quantities detected in THP-1 cell lysates at 72 h post infection.

**Table 2.**
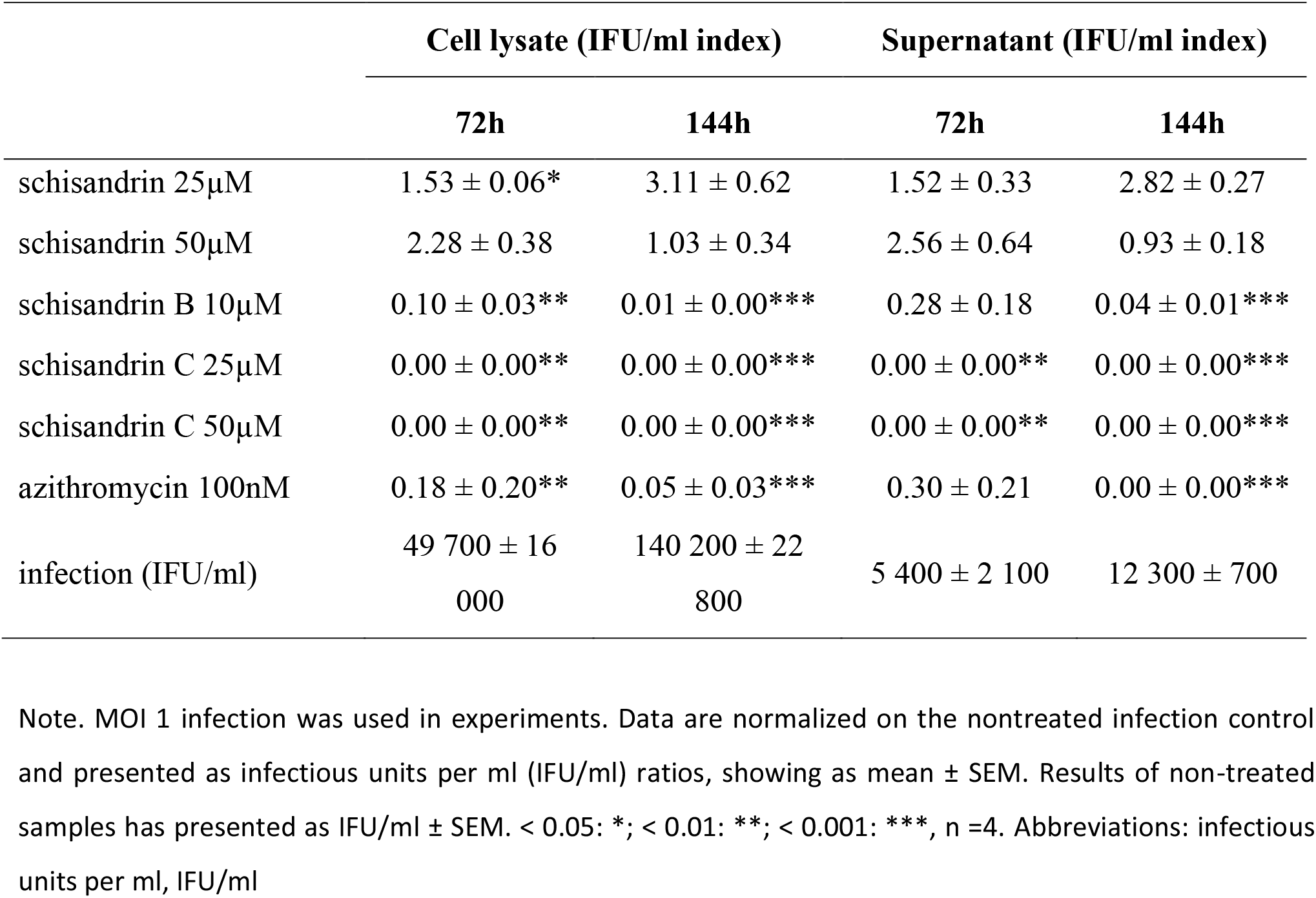
The Impact of schisandrin lignans on *C. pneumoniae* infectious progeny production.

### Role of cellular redox status in the effects of schisandrin lignans on C. pneumoniae infection

As key players in innate immunity, macrophages respond to microbes and other danger signals by generating effector molecules such as reactive oxygen species (ROS) and nitric oxide (NO) intended to kill the pathogen ^30^. As the dibenzocyclooctadiene lignans are known for their redox activities and changes in cellular redox status have also been linked to bacterial persistence, we extended the study on differential effects of schisandrin and schisandrin C on *C. pneumoniae* infection by addressing changes in cellular redox status. As shown in Fig. 3A, both MOI1 and MOI5 *C. pneumoniae* infections elevated the ROS production of the infected macrophages statistically significantly at 48 h post infection. After 72 h infection ROS levels were elevated in MOI5-infected samples. As also total cellular ATP levels were elevated in the infected samples (supplementary fig. 1) but no increase in NADPH oxidase activity was observed (data not shown), this may reflect the manipulation of host cell mitochondrial function in a manner similar to a related pathogen *C. trachomatis* ^31^. While NADPH oxidase is considered the major source of ROS in stimulated macrophages in general, previous studies have proposed that mitochondrial ROS production also takes part in macrophage responses to bacterial invaders ^32^. In macrophages, the mitochondrial ROS have been reported to actually contribute to chlamydial survival via mechanisms involving NLRP3 inflammasome activation ^33, 34^.

**Figure 3.**
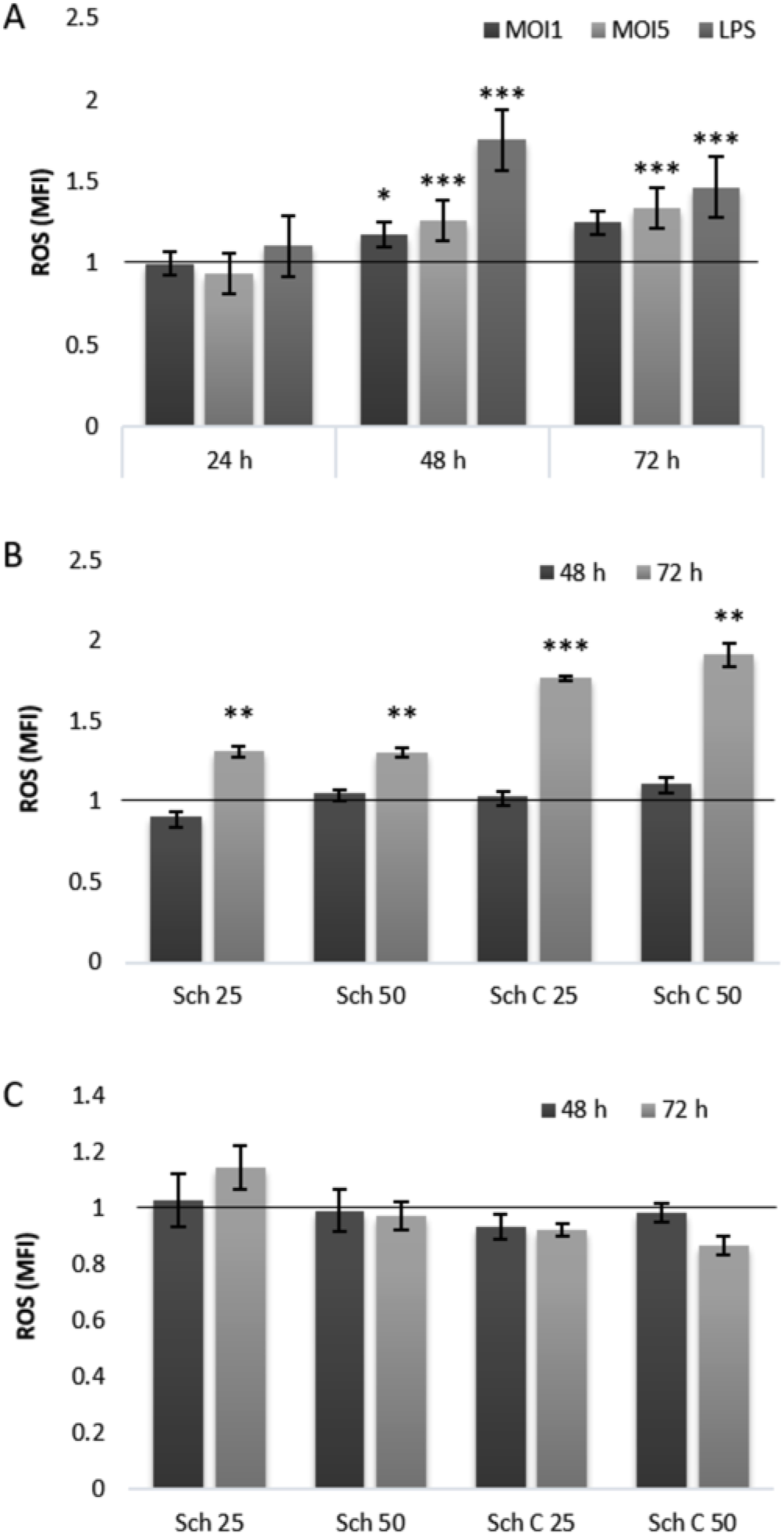
Impact of *C. pneumoniae* infection and schisandrin lignans on intracellular ROS production. (A) THP-1 macrophages were infected with *C. pneumoniae* at MOI1 and MOI5 or treated with 1μg/ml of LPS for 24, 48 or 72 h. (B) THP-1 macrophages were exposed to schisandrin lignans for 48 and 72 h. (C) THP-1 cells were simultaneously infected with *C. pneumoniae* and treated with schisandrin lignans for 48 and 72 h. Intracellular ROS levels were measured after 30 min incubation with DCFH-DA and followed by 3 h incubation with PBS. Data are normalized as a ratio of 0.25 % DMSO cell control (A, B) or infection control (C) and shown as a mean ± SEM. Statistical significance is presented as marks of P values: < 0.05: *; < 0.01: **; <0.001: ***, n=4. Abbreviations: Reactive oxygen species, ROS; Mean fluorescence intensity, MFI; Multiplicity of infection, MOI; lipopolysaccharide, LPS; Schisandrin, Sch.

As the data presented in Fig. 3 confirmed the relevance of ROS within *C. pneumoniae* infection in THP-1 macrophages, we evaluated the impact of schisandrin and schisandrin C in this respect. No differences in basal ROS levels were observed after 4 – 48 h exposure, but after 72 h schisandrin and schisandrin C elevated ROS levels at both 25 μM and 50 μM concentration (Fig. 3B). The concomitant administration of schisandrin and schisandrin C with MOI5 *C. pneumoniae* infection did not change the detected ROS levels compared to a vehicle-treated control infection (Fig. 3C).

Earlier work on anti-persister agents has indicated that promotion of oxidative stress by increased rOS production eradicates persister bacteria and enhances bacterial killing by conventional antibiotics ^35, 36^. Despite the impact of the lignans on macrophage basal rOS levels, ROS promotion is not likely the primary mode of action of the compounds against *C. pneumoniae*, as the infection-induced ROS levels show no difference between treated and non-treated cells. Furthermore, similar impact on basal ROS levels was observed for schisandrin and schisandrin C despite their opposite effects on *C. pneumoniae* progeny yields (Table 2).

The impact of *C. pneumoniae* infection and schisandrin lignans on redox status of THP-1 macrophages was further studied with determining cellular GSH concentrations after infection, lignan treatment, or the combination of these two. As shown in Fig. 4A, *C. pneumoniae* infection caused a time- and infection MOI-dependent elevation in cellular GSH levels, detectable 48 – 72 h post infection. Time-dependent fluctuation of GSH pools after chlamydial infection have also been reported by others ^37, 38^ and can be considered a homeostatic response of the host to the elevated ROS levels.

**Figure 4.**
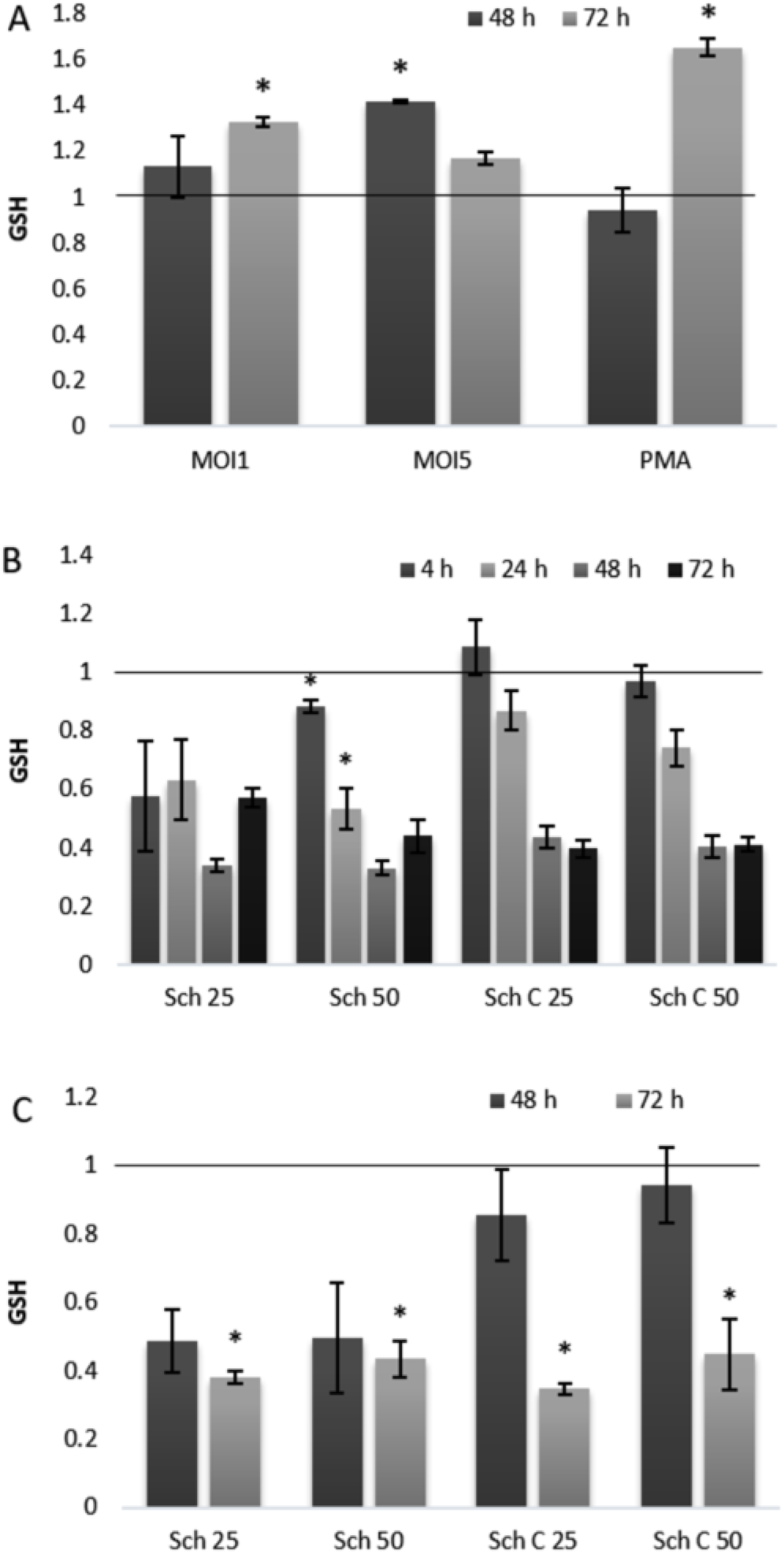
Effects of schisandrin lignans on total cellular GSH levels in THP-1 macrophages. (A) THP-1 cells were infected with *C. pneumoniae* at MOI1 and MOI5 for 48 and 72 h and PMA was used as control. (B) THP-1 cells were exposed to schisandra lignans for 4 to 72 h. (C) Cells were infected with MOI5 and exposed to schisandra lignans for 48 and 72 h. Total basal GSH concentrations were determined and normalized to total protein concentration of the sample. Data are shown as a ratio of 0.25 % DMSO control and shown as mean ± SEM. Statistical significance is presented as marks of P values: < 0.05: *; < 0.01: **; < 0.001: ***. N=4. Abbreviations: Glutathione, GSH; Multiplicity of infection, MOI; phorbol-12-myristate-13-acetate, PMA; Schisandrin, Sch.

Our replication results show that despite the elevated ROS levels (Fig. 3A) and altering GSH levels (Fig. 4A) during infection, a significant fraction of the bacteria maintains an actively replicating phenotype. Thus, in contrast to murine macrophages ^17, 39^ oxidative stress seems not to result in purely persistent infection phenotype in THP-1 macrophages.

Previous work has highlighted the role of NO and inducible nitric oxide synthase (iNOS) as a trigger for chlamydial persistence in murine RAW264.7 macrophages ^17, 37^. NO production in in PBMCs and macrophages involves, however, remarkable inter-species differences ^40, 41^ and based on data published by us and other research groups, the human THP-1 cell line produces NO neither in its monocytic nor macrophage like phenotype ^42–44^. The lack of iNOS induction after chlamydial trigger can be speculated to enable the productive *C. pneumoniae* infection in THP-1 macrophages, yet more comprehensive picture on host cell redox status is necessary in order to draw conclusions in this respect.

The medicinal plant-derived schisandrin lignans have recently been reported to harbour pharmacological activities, such as neuro- and cytoprotective as well as anti-inflammatory properties ^45^. A key mechanism mediating these activities is considered to be the lignans’ impact on cellular redox status. The promotion of redox activities is linked to their ability to modulate mitochondrial functions ^46, 47^. In addition, our previous studies in monocytic THP-1 cells showed that the lignans affect cellular glutathione metabolism by causing a decrease in total GSH pools ^42^. Similar to monocytic THP-1 cells, THP-1 macrophages exhibit remarkably lowered GSH pools after lignan treatment (Fig. 4B), and a drastic decrease in GSH levels was also observed in *C. pneumoniae*-infected THP-1 macrophages after lignan exposure (fig. 4C). The relevance of GSH depletion in the lignans’ activities on *C. pneumoniae* was confirmed by supplementing the infected cultures with GSH ethyl ester. While administration of this cell-permeable GSH derivative alone increased EB yields by approximately 40 %, supplementation of schisandrin-treated infections yielded infectious progeny levels similar or lower than those in the infection control, indicating that the GSH supplementation eliminates the elevating effect of the 25 μM schisandrin on bacterial progeny production.

The cellular GSH balance may affect intracellular bacteria by a variety of mechanisms. GSH has been found to act as a major cysteine source of intracellular bacteria ^48^ and it has been reported to indirectly affect chlamydial energy supply by increasing cell wall permeability ^49^. GSH depletion is also known to induce K+ efflux ^50^ which, in turn, can promote the chlamydial replication via the induction of NLRP3 inflammasome in the host cells ^33, 51^. On the other hand, GSH and its metabolites are directly toxic to some intracellular bacteria ^52^ and the GSH-dependent changes in cellular redox status may detrimentally affect *C. pneumoniae* survival, as suggested in murine models ^37^. Interestingly, GSH is also known to induce virulence gene expression of *Listeria monocytogenes*^53^, implying that it may serve as an indicator of the local environment, directing virulence gene expression of intracellular bacteria. To date, processes linked to the ability of *Chlamydia* spp. Bacteria to sense their microenvironment have remained poorly understood, and the potential role of GSH in chlamydial adaptation and balance between active and persistent phenotype warrants further investigation.

Despite their similar effects on cellular ROS and GSH levels, schisandrin and schisandrin C show differential activity on the *C. pneumoniae* infection in THP-1 macrophages. This may reflect two separate aspects of their biological activities: a redox-dependent phenotypic switch by *C. pneumoniae* from persister to active replication and a dual mode of antichlamydial action by schisandrin C. While schisandrin C exhibits chlamydiocidal activity in the acute infection model, schisandrin does not affect actively dividing bacteria at 25 μM concentration ^29^. Based on these observations, we propose that depleting cellular GSH stimulates chlamydial growth, and persistent infection is converted to active state. In the case of schisandrin this is seen as the promotion of infectious progeny formation, whereas schisandrin C acts as a multimodal antibacterial agent yielding a potent eradication of macrophages by killing the bacteria it is switching towards active replication.

To date, only a few drug-like molecules have been described as phenotypic switchers reverting persister bacteria from dormancy to active growth ^2, 54, 55^ and to our knowledge, the schisandrin lignans represent the first phenotypic switchers described to be active on intracellular bacteria. Based on rodent bioavailability studies, micromolar plasma concentrations of the lignans can be achieved after a single oral dose ^56^, indicating the potential of the lignans as leads for orally administered drugs. Furthermore, our in vitro cell viability data (Supplementary information, ^29, 42^ and the numerous in vivo studies on these compounds ^45^ indicate the lack of acute or subacute toxicity of the lignans.

To conclude, the constant increase of detected number of bacterial copy numbers and infectious progeny in our studies indicates that *C. pneumoniae* is able to establish a productive infection in THP-1 monocyte-derived macrophages. The limited potency of azithromycin in clearing the cultures from bacteria implies to the presence of an antibiotic-tolerant subpopulation of bacterial and emphasizes the importance of including nonpermissive host cell lines in chlamydial susceptibility studies. Redox status and glutathione homeostasis have recently emerged as modulators for intracellular bacteria virulence ^53, 57^, indicating that targeting cellular redox mechanisms may offer a means for inducing a phenotypic switch in pathogenic bacteria. The superiority of schisandrin C compared to azithromycin in its ability to eliminate both active and persistent bacteria in THP-1 macrophages highlights the potential of this strategy in anti-persister therapy, triggering future research on nonconventional antibacterials.

## Acknowledgements

The authors wish to thank the funding bodies for the financial support.

## Transparency declaration

The authors declare no conflicts of interest.

